# Bioelectronic Zeitgebers: targeted neuromodulation to re-establish circadian rhythms

**DOI:** 10.1101/2023.04.30.538861

**Authors:** Alceste Deli, Mayela Zamora, John E. Fleming, Amir Divanbeighi Zand, Moaad Benjaber, Alexander L. Green, Timothy Denison

## Abstract

Existing neurostimulation systems implanted for the treatment of neurodegenerative disorders generally deliver invariable therapy parameters, regardless of phase of the sleep/wake cycle. However, there is considerable evidence that brain activity in these conditions varies according to this cycle, with discrete patterns of dysfunction linked to loss of circadian rhythmicity, worse clinical outcomes and impaired patient quality of life. We present a targeted concept of circadian neuromodulation using a novel device platform. This system utilises stimulation of circuits important in sleep and wake regulation, delivering bioelectronic cues (Zeitgebers) aimed at entraining rhythms to more physiological patterns in a personalised and fully configurable manner. Preliminary evidence from its first use in a clinical trial setting, with brainstem arousal circuits as a surgical target, further supports its promising impact on sleep/wake pathology. Data included in this paper highlight its versatility and effectiveness on two different patient phenotypes. In addition to exploring acute and long-term electrophysiological and behavioural effects, we also discuss current caveats and future feature improvements of our proposed system, as well as its potential applicability in modifying disease progression in future therapies.

## I. Introduction

Disordered sleep and wakefulness are secondary features of neurodegenerative conditions. These features severely impact patients’ daily functioning and overall quality of life, and can range from sleep fragmentation and excessive daytime sleepiness (EDS) to profound loss of circadian rhythmicity, as observed in advanced delirium and dementia [1]. Underlying this behavioural dysregulation are pathological patterns of electrophysiological activity that correlate with the sleep-wake cycle. As part of the aging process, cortical power spectra shift towards faster frequencies during sleep. This is evidenced by reduced delta-band (0 - 4 Hz) power, less time spent in slow-wave sleep, and increased sleep fragmentation [2]. This phenomenon is aggravated in neurodegenerative disorders [3], [4]. In addition, during neurodegeneration wake rhythms are often characterised by electroencephalogram (EEG) slowing – a reverse pattern that is associated with worse clinical outcomes [5], [6].

These pathological trends in electrophysiological patterns during sleep and wakefulness, arising from a variety of primary pathologies, such as tau aggregation in Alzheimer’s Disease (AD) or alpha-synuclein accumulation in Multiple Systems Atrophy (MSA), all point towards dysfunction of fundamental processes that underlie cortical dynamics. From a network perspective, a group of subcortical nuclei such as those comprised of the reticular activation system (RAS) provide ascending projections to key cortical regions important for wakefulness [7]. Brainstem arousal networks that pertain to the RAS are one of the earliest sites of neurodegenerative changes, well before the development of overt disease-specific symptomatology [8]. Moreover, many of these regions physiologically exhibit diurnal firing patterns [9].

Current surgical management of neurodegenerative disorders via surgical neurostimulation (a treatment option for pharmacoresistant neurodegenerative conditions) imposes constant stimulation settings that are agnostic to diurnal and circadian patterns of brain activity. Most brain regions chosen for functional neurosurgical targeting are selected based on motor function modulation – these include areas with dual roles in arousal regulation as well as other functions. The use of constant deep brain stimulation (DBS) settings may therefore influence circadian rhythmicity and aggravate sleep disruption [10]. For these cases, tailored DBS settings aligned to the sleep/wake cycle may be more appropriate for delivering therapy. Moreover, fine-tuning DBS therapy in this manner to enable potential entrainment or reinforcement of physiological oscillatory patterns associated with wakefulness and sleep may lead to improved therapeutic outcomes.

In this paper we explore the concept of delivering DBS therapy aligned with the sleep-wake cycle, with the goal of reinforcing physiological oscillatory patterns associated with sleep and wakefulness. Tailored and scaleable neurostimulation cues, related to sleep/wake neuronal processes, are what we will refer to as ‘bioelectronic Zeitgebers.’ These cues, delivered by our novel programmable neurostimulation device to neuronal nodes important for arousal modulation, can be carefully selected and timed to achieve the desired impact on sleep and wakefulness. In support of this concept, we also present preliminary evidence of the approach’s capacity to modify neurophysiology (evidenced by variations in day- and night-time recorded electrophysiological activity) and behaviour (such as changes in subjective EDS ratings and patient-reported nap diaries). We propose that aligning DBS therapy with the sleep-wake cycle in this manner may provide improved symptom control in other specific disease phenotypes, in addition to potentially impacting longitudinal disease progression.

## II. Methods

We will present our key elements of deployment of ‘bioelectronic Zeitgebers’ through an experimental device (currently embodied in the Picostim-DyneuMo), in order to ameliorate sleep-wake pathology and improve quality of life in our patient populations. We will therefore explore factors that may facilitate generalisation to other conditions and/or target nuclei, as well as more detailed algorithm parameter selection and trial- or target-specific methodology.

### A. Brief device specification: Picostim–DyNeuMo Overview

The Picostim–DyNeuMo research system is a cranially-mounted, rechargeable neuromodulation device that enables dynamic variation of delivered stimulation programs in response to movement (Mk-1) [11], time (Mk-1 and -2) [12], and sensed biopotential signals (Mk-2) [13]. These embedded device capabilities are enabled via an onboard device accelerometer, clock and filters which enable configurable algorithm deployment. This device therefore offers the capability for tailored diurnal and circadian algorithm creation and use in a variety of neurological indications where diurnal and circadian symptom modulation may be an aim.

### B. Network Node Selection: Targets for Sleep/Wake Modulation

There is an array of both extrathalamic (brainstem/RAS) and thalamic brain regions (such as central thalamic nuclei supporting forebrain mesocircuit function [14]) whose modulation may affect sleep and wakefulness. The choice of implantation target relies on both underlying condition and pathophysiology. For instance, where circadian or sleep disorder is a neurodegenerative sequela in the presence of combined motor and autonomic dysfunction, a brainstem RAS target could be chosen to ameliorate additional pathology. In another example, in the absence of motor pathology but where cognitive deficits and/or a global decrease in level of arousal are predominant, thalamic regions could also be selected [15]. However, the presence of severe thalamic pathway damage (as in disorders of consciousness following traumatic brain injury) could favour an RAS target with independent cortical projections. The neuromodulatory technology that we propose could therefore be deployed via a variety of ‘nodes’ in networks modulating sleep and wakefulness, depending on the underlying condition and network availability.

The implantation site for the first use of such a bioelectronic research system was the pedunculopontine nucleus (PPN) (Fig. 1) (as part of the MINDS clinical trial, ClinicalTrials.gov Study Identifier: NCT05197816). The PPN is part of brainstem arousal circuits that deliver wake-related signals to cortex, both through thalamic relay nuclei (parafascicular nucleus thalami) and through an extra-thalamic pathway [16]. Preferentially active during wakefulness and REM sleep, its reported firing rates lie within the beta/gamma range (20-60Hz) with current clinical stimulation settings at 40 Hz often said to reflect a physiologic wake frequency [17], [18]. In keeping with its important role in sleep/wake activity, findings from our group have also shown that slow-wave sleep can be affected by PPN DBS timing [19]. The PPN additionally has long-range connections with areas important for gait and locomotion (notably basal ganglia) [20]. Its degeneration has been linked to both motor and non-motor symptoms in neurodegenerative disorders, while its stimulation has also been reported to affect locomotion and autonomic function [21], [22].

**Fig. 1.**
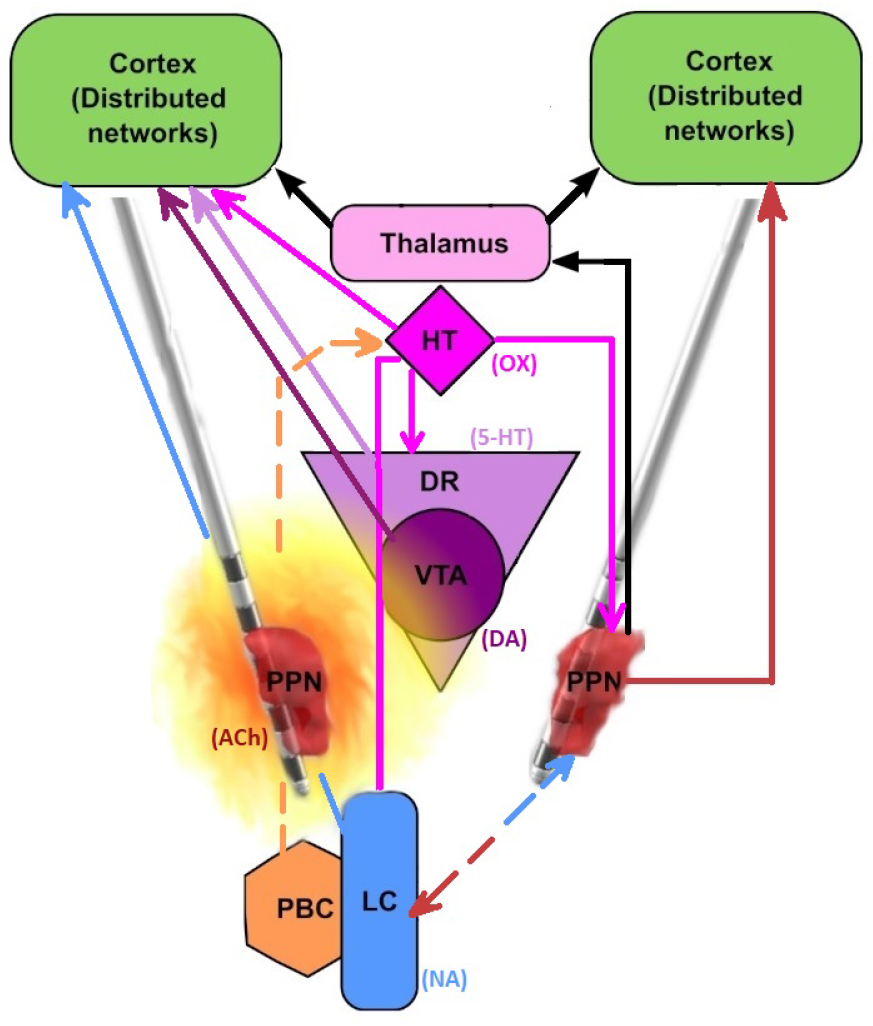
The PPN is located in a close-knit, spatially distributed complex network of subcortical areas and neurotransmitters, regulating sleep and wakefulness. The effect of circadian therapy may be a result of interface with either its own projections (simplified, right side) or as a function of volume of DBS current-activated tissue, given that brainstem nuclei are very closely situated. Activated tissue is not a homogeneous sphere and may include projections from subcortical nuclei (such as the locus coeruleus, LC) to cortex. Note that network projections may be bi-directional (LC-PPN), while some regions can have global regulatory roles (such as hypothalamic-HT areas with orexinergic-OX projections). OX: orexin, 5-HT: serotonin, DA: dopamine, NA: norepinephrine, ACh: acetylcholine. Other nuclei listed: PBC: parabracial nuclei complex. DR: dorsal raphe nuclei. VTA: ventral tegmental area. HT: hypothalamus.The PPN is visualised here in darker red, as mapped using imaging analyses in an MSA patient (LeadDBS software).

### C. Sleep-Wake Cycle Aligned Stimulation: Algorithm Overview

The embedded, configurable algorithm that we propose and have tested was built on the principles of bioelectronic Zeitgeber therapy, responsive to temporal activity patterns and delivered through implantable devices in the aforementioned groups of brain nuclei. Therefore, it consists of three main layers of control for adapting stimulation parameters to daytime and night-time operation: (1) diurnal adjustments to the basal stimulation frequency and amplitude (broadly differentiating wake and sleep), (2) motion-adaptive updates to the stimulation frequency and amplitude, which can for instance be deployed during night-time operation (Fig. 2) and (3) state-specific adjustments (such as discrete adjustments per sleep stage), aimed at more tailored protection of neural function within the broader concepts of sleep and wakefulness. To illustrate each example, we will describe how the Picostim-DyNeuMo was configured to deliver day- and night-time stimulation programs to reinforce oscillatory activity of the PPN, associated with each phase of the sleep-wake cycle.

**Fig. 2.**
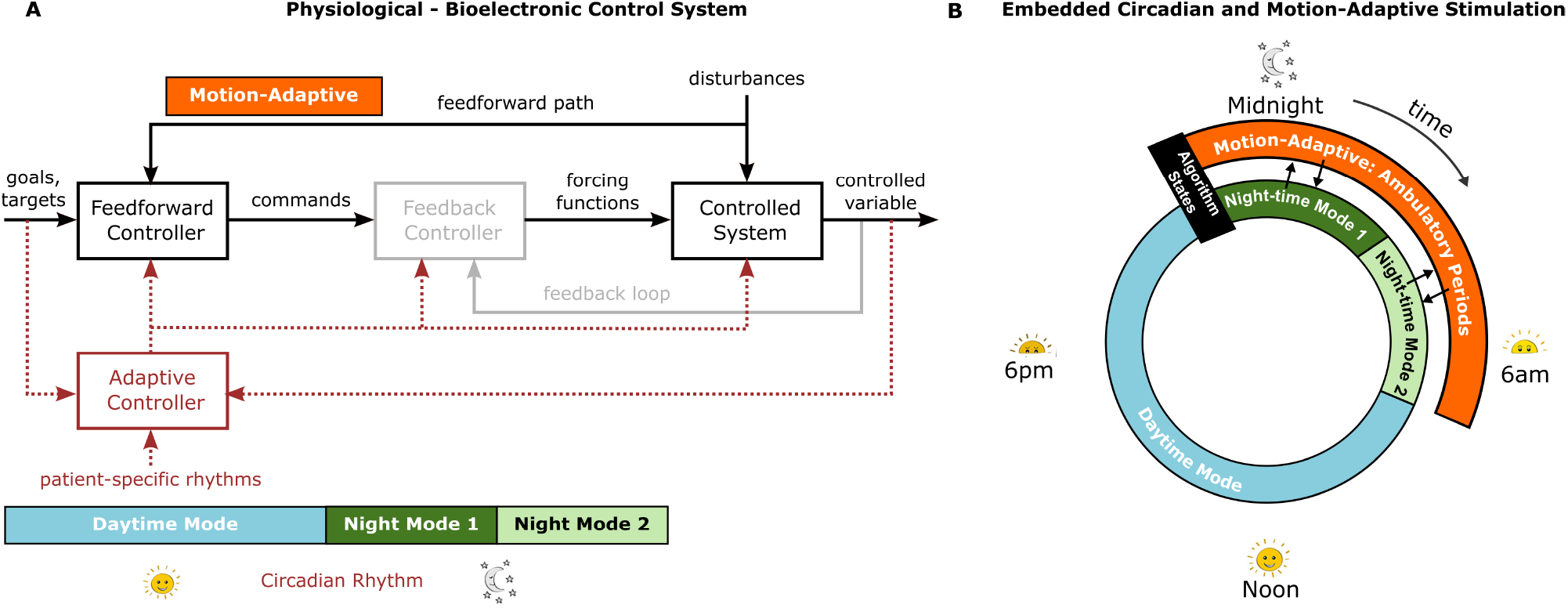
Bioelectronic Zeitgeber control strategy using a combination of scheduled and motion-adaptive stimulation to entrain physiological, circadian patterns. A: Block Diagram of the control structure for the physiological system. Stimulation parameters are based on patient-specific rhythm timings that can be further tailored to pathology and ongoing neural activity (adapted from [23]). Implementation of the feedback controller block (grey) proposed in [23] is not included in this present study. The output of the feedforward controller goes directly to the controlled (physiological) system. B: Illustration of the embedded circadian and motion-adaptive stimulation algorithm for adaptation of stimulation parameters over the 24 hour sleep-wake cycle.

Sustained wakefulness of patients during daytime does not only translate to a higher level of engagement with their carers and surroundings. Fewer unplanned bouts of daytime sleep translate to greater buildup of homeostatic sleep pressure, priming patients for earlier sleep onset, shorter slow-wave sleep latency and better sleep quality at night. To discourage bouts of drowsiness and unplanned daytime sleep, daytime DBS settings should be established to entrain cortical rhythms towards higher frequencies and avoid EEG slowing through activation of arousal-promoting pathways. In our trial in the PPN, this was implemented by utilizing constant gamma frequency stimulation during the daytime (such as at a 40Hz stimulation frequency), while normalised PPN activity may also support healthy motor patterns. Where that is the case, enhanced physical activity may become an additional ‘Zeitgeber’ reinforcing wakefulness.

Motion-adaptive updates to stimulation settings may be particularly important in the night-time, when patients may require assistance in order to be able to mobilise (and tend to their needs, since control of bodily functions may be affected by motor dysfunction or dysautonomia). In our illustration of how the Zeitgeber strategy is implemented in the Picostim-DyNeuMo (Fig. 2), the outer ring of stimulation illustrates motion-adaptive adjustments to the stimulation amplitude and frequency, in order to facilitate night-time ambulatory periods. When motion or postural change exceeding a set threshold is detected, the stimulation amplitude and frequency are updated to wake-related settings, to avoid potential postural drops in blood pressure and alleviate disease-related motor symptoms. Stimulation is maintained at these parameter values until the next scheduled adjustment from the circadian scheduler, which reverts the stimulation parameters to their corresponding circadian-scheduled values.

With regards to DBS night-time settings, tailored stimulation cues delivered at the neuronal nodes of arousal-promoting networks should respect the physiological patterns of oscillatory activity that are observed during sleep, as well as promote a normalisation of sleep architecture where there is stagespecific pathology. In our PPN example, we programmed the Picostim-DyNeuMo in order to mimic the region’s nocturnal quiescence (especially during slow-wave sleep), therefore avoiding daytime amplitudes in gamma settings. Furthermore, we did not select a gamma stimulation frequency during the first half of the night but instead, opted for lower frequency settings -especially important in the case where there was specific REM-sleep related pathology, usually occurring early during the night. In Fig. 2 this is showcased as different settings in the first half of the night versus the second half of the night (normally, REM sleep periods become more frequent towards the morning). We will be further examining the capacity for sleep stage-specific enhancement in future patients and across the scope of our collaborative work.

Another, longer-term level of Zeitgeber fine-tuning would rely on monitoring for circadian phase shifts, such as increasingly earlier wake-up and bed times across weeks and months, a phenomenon termed phase-advancement. These could occur either as response to therapy or as a function of disease progression. A progressive normalisation of chaotic sleep/wake behaviours, which are present in both neurodegenerative conditions and mental health co-morbidities, would mean that settings would also need re-adjustment to the emerging patterns. This gains further importance if we hypothesise that effects on circuit dynamics may also be consolidated through stimulation-related plasticity changes of the underlying arousal-promoting networks.

### D. Trial-specific and Data-gathering Concepts

We will follow the description of generalisable algorithm methods with some details specific to the MINDS trial, especially with regards to its elements relevant to sleep/wake and circadian function modulation. This is a clinical feasibility study, investigating adaptive DBS for MSA, a severe neurodegenerative condition characterized by both debilitating autonomic, postural and gait impairments in addition to the presence of sleep/wake pathology [24]. Therefore the target selection (PPN) has been planned in order to ameliorate a constellation of motor and non-motor symptoms.

To capture clinically relevant aspects of disrupted sleep/wake cycles, a combination of behavioural assessments and electrophysiological monitoring is utilised. Upon enrollment, a dedicated set of validated questionnaires is administered, in addition to other disorder-specific symptom assessments. These questionnaires center on EDS (captured by the Epworth Sleepiness Scale, ESS [25]), subjective sleep quality and associated metrics (Parkinson’s Disease Sleep Scale PDSS [26]). The presence of REM sleep-related pathology, common in MSA, is also assessed (REM Sleep Behaviour Disorder Screening Questionnaire, RBDSQ [27]). Baseline wake and sleep EEG (polysomnography: a combination of EEG, eye movement monitoring and muscle tone) as well as continuous electrocardiogram (ECG) are obtained at baseline and upon recovery post-implantation. With regards to wake states captured, parameters such as eye closure, medication status and time-of-day are controlled for between recording sessions, to account for baseline variation effects.

The same assessments are obtained upon switch-on of the implanted device and at set intervals longitudinally thereafter, accompanied by patient sleep and nap diaries (completed by the carer-observer). Subjective patient metrics are complemented by neurophysiological measurements which are obtained during planned visits over a six-month course, to ensure the fine-tuning of stimulation parameters and related algorithms. Once the tailor-made night-time algorithm has been finalized, its effects on sleep physiology are then compared to a (control) night of continuous daytime stimulation (as recorded by polysomnography - PSG). This combination of objective and subjective measures ensures both biomarker tracking and correlation to patients’ behavioural outcomes, experiences and perceived quality of life.

Electrophysiological (both daytime and PSG) data collection is subject to trial-specific adaptations related to the implantation process for the Picostim-DyNeuMo in this specific target region. To ensure the reproducibility of anatomical location, given the complexity of brainstem anatomy, in addition to localization steps reported previously by our group [28], pre-operative planning involves a dedicated mask, based on a neuroimaging atlas of the relevant ascending arousal circuits [29]. Another operative feature due to the cranially-mounted device is the creation of a frontocentral skull pocket, which houses the device implantable pulse generator (IPG). The cranial placement of the IPG means that EEG sampling of cortical activity may slightly vary in central regions between patients. This is a consideration that subsequently affects the recording electrode montages selected during longitudinal patient monitoring.

### E. Data Analysis

Wake electrophysiological data was pre-processed in Matlab (2022a, Mathworks Inc., Nantick, MA, USA), using a set of 3rd order Butterworth filters (high-pass: 1 Hz and lowpass: 130 Hz), a band-stop filter to avoid line noise and its harmonics, as well as de-trending during visualisation to eliminate any drifts. With regards to polysomnography data, these were pre-processed and analysed as per American Academy of Sleep Medicine (AASM) guidelines [30]. Behavioural data was analysed using two-sample t-tests as well as ANOVA (with Bonferroni correction for multiple comparisons).

## III. Results

In support of this bioelectronic Zeitgeber concept, we present data from two patients where the system has now been implanted and activated, and which have been discharged from hospital directly to their natural environment. These illustrate how the concept can be generally applicable in two discrete complaints relevant to sleep/wake pathology, as well as how the algorithm can undergo individual tailoring in order to ameliorate patient-specific symptomatology. Specifically, one case presented with a known diagnosis of REM sleep behavior disorder (persisting despite pharmacological treatment), however excessive daytime sleepiness (EDS) was not a major complaint. In the other case, although there was no formal diagnosis of sleep disorder, a significant amount of EDS was present. Written informed consent was obtained in both cases upon enrollment and pre-implantation, while the procedure and subsequent inpatient care was in compliance with local Ethics and the Declaration of Helsinki. Consent was also verbally revisited prior to all recordings and non-invasive trial procedures.

### A. Zeitgebers of RAS Stimulation: Daytime reinforcing of wake-related cortical activity

Along the lines of our described bioelectronic Zeitgeber strategy, we have preliminary proof of broad cortical circuit engagement during daytime PPN gamma stimulation from both patients. Acute effects of daytime parameters on the power spectra involve non-linear acceleration of peak beta frequency (13-30 Hz) in response to stimulation, when compared to an off-stimulation state within each testing session in both patients (Fig. 3A). The maximum of these effects appeared adjacent to the half-harmonic of the applied gamma DBS frequency, further evidenced by peri-harmonic shifts depending on stimulation frequency that is applied within the same experimental session (Fig. 3B.). Daytime beta oscillations in frontal and prefrontal areas have been linked with top-down control of behaviour and cognitive function [31]. There is emerging evidence in our data that beta oscillatory dynamics in these cortical regions may be chronically impacted by daytime RAS stimulation, even in an off-stimulation state, further supporting the claim that DBS is engaging key projections to cortex. Follow-up data obtained for the same prefrontal cortical brain area, time-of-day and behavioural state suggest that longitudinal effects of stimulation settings on cortical activity include shifts of the existing maximal frequency content towards higher values in that band and the half-harmonic of the stimulation frequency, as well as an overall increase in beta frequency content (Fig. 4). The quantification of such results and the formal exploration of their relationship to the concept of entrainment is ongoing.

**Fig. 3.**
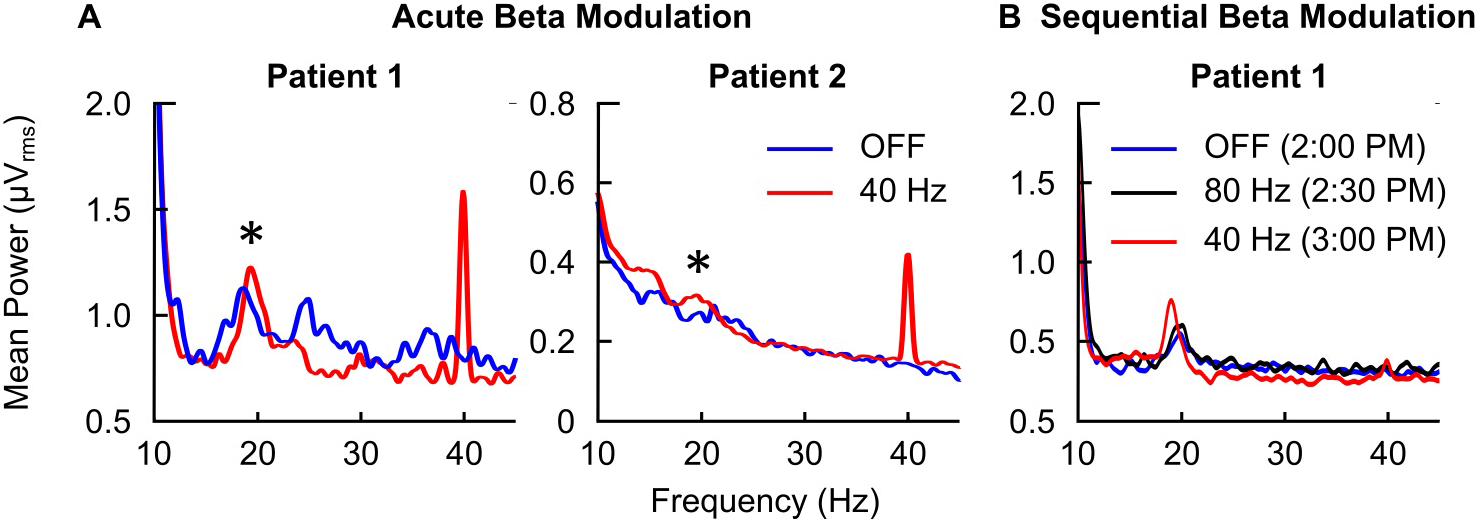
Modulation of network neurophysiology, as recorded by wake EEG during different time points and stimulation settings in both patients. A: Acute beta modulation within-patient and testing session (wake, resting state with eyes closed). Linear power spectral plots reflecting modulation in beta (13-30 Hz) frequency content during PPN gamma stimulation at 40Hz. Changes are observed during the stimulation half-harmonic (20Hz), as highlighted by asterisks overlaying that frequency in the sub-plots. B: Further evidence of a relationship with the stimulation half-harmonic is offered when sequential changes in DBS frequency are applied. Sample data from two patients (‘Patient 1’and ‘Patient 2’) are presented.Identical recording montages (fronto-occipital) have been used for between-patient comparisons. A more conservative (less spatially distributed) montage was used for sub-plot B, hence the presence of the stimulation peak at 40Hz is less obvious.

**Fig. 4.**
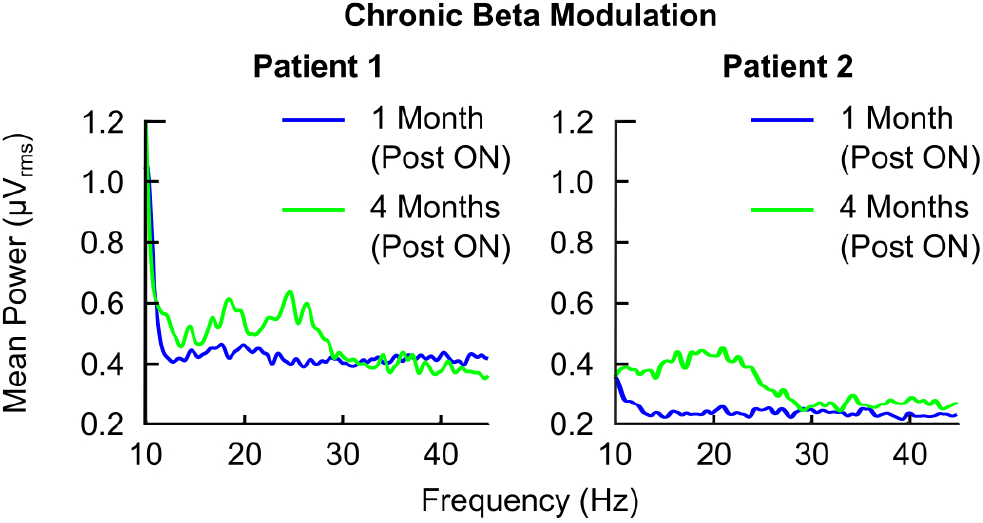
Chronic modulation of circuit neurophysiology. Evidence of chronic cortical circuit modulation by RAS/PPN DBS in both patients. Intrinsic beta content during the ‘OFF’ stimulation state is increased over time in both cases (frontal EEG channels, same re-referencing montage). Recordings were made at the same time-of-day within-patient, during both patients’ follow-up sessions. Behavioural recording conditions as well as the patient’s medication state were also constant.

### B. Phenotype-specific effects: Results on Excessive Daytime Sleepiness (EDS)

In addition to general effects on diurnal physiology, which could also improve longer term daytime cognitive functioning, we further illustrate how tailored adjustments in monitoring and programming can improve specific behavioural patterns and therefore may directly the quality of life of patients with specific sleep/wake complaints. In the EDS phenotype, diurnal bioelectronic Zeitgeber delivery reduced bouts of daytime sleepiness, as reflected on both subjective reports and observer daily diary entries over a period of 15 weeks. A subjective decrease in overall time spent asleep during the day was reported acutely after switch on, as also indicated by a normalisation of Epworth scores (which reflect the perceived degree of sleepiness in a variety of social situations by the patient). This was confirmed through our analyses, with the second week compared to the first having a significant reduction in disruptive daytime sleep duration (p = 0.0022), with a mean reduction of circa an hour of daytime sleep (61.9 minutes, CI 11.8 to 112.6 minutes, Fig. 5B). We also noted that, compared to the first week of stimulation, time of the last nap occurred significantly earlier over time (by around 5 hrs) (Fig. 5A) (p = 0.0036 with mean difference of 5.3 hours and CI from 0.9 to 9.7 hours, between weeks 1 and 12).

**Fig. 5.**
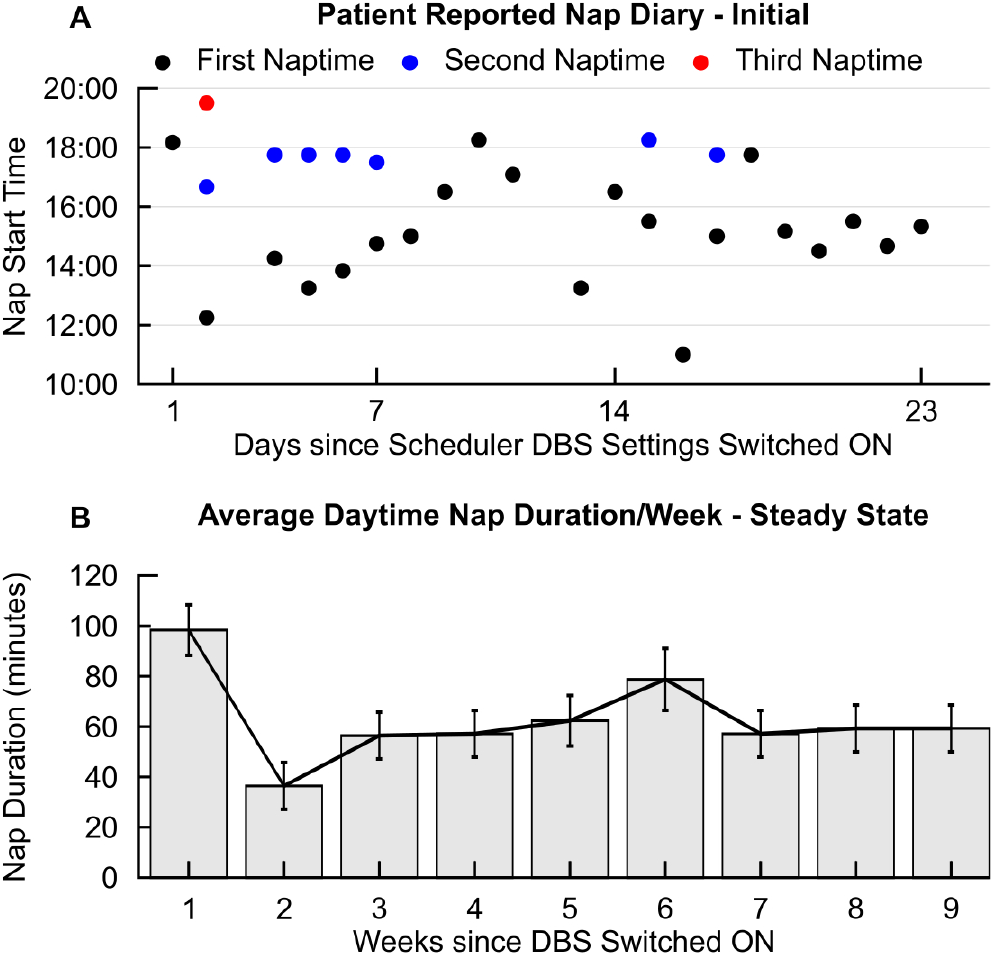
Modulation of symptomatology in the MSA phenotype presenting with Excessive Daytime Sleepiness (EDS). A: Shift to earlier nap times and reduced number of naps across days from device activation (switching ‘ON’ of scheduled settings), in the EDS phenotype. B: Reduction of total average nap duration (minutes) across days per week, from first week after device activation (switching ON). Device activation took place three months post-implantation, ensuring potential acute post-operative effects (such as oedema-related changes and stun effect) had resolved and recovery achieved. We noted a transient response with an initial acute reduction in nap duration (week 1 to week 2), while over time for this contact combination there seems to be a ‘steady state’ (as evidenced by the plateau from weeks 7 to 9).

During programming in the EDS phenotype, we focused on monitoring daytime cortical frequencies and opted for a scheduled distribution of settings which maximised alertness-while ensuring that motor and autonomic function, as well as sleep integrity, were not directly impaired during changes to scheduled settings. Despite an overall deterioration in terms of disease progression, this observed reduction in disruptive daytime sleep duration was longitudinally maintained over a period of 15 weeks (week 15 had an average of 51.2 minutes spent in daytime naps compared to week 1, with p = 0.0386 and wider CI however, ranging from 1.04 to 101.34 minutes). We finally saw an element of longitudinal reduction in nap number, from 2 naps as a daily average during the first week to 0.86 naps per day in week 15 (p = 0.0279 for a reduction in 1.14 nap and CI between 0.05 and 2.35).

### C. Phenotype-specific effects: Results on Disordered and Fragmented Sleep

We will also showcase some representative data from our patient with a formal diagnosis of sleep disorder (REM Sleep Behaviour disorder, RBD), illustrating the patient-specific need for stimulation adjustments according to sleep stage pathology, in order to obtain optimal results. This diagnosis is characterised by loss of muscle atonia during REM, vivid dream re-enactment and associated sleep fragmentation as a consequence of high arousability, which are consciously adversely experienced by patients and reliably reportable in questionnaire-based diagnostic screening [28]. Initial nighttime settings tailoring in the RBD phenotype included reports of frequent and early arousals, as well as vivid dream content. Stimulation settings were therefore adjusted with repeat RBD screening confirming that there was no clinical risk of disordered REM sleep. These reports may reflect stimulation-induced PPN hyperactivity consistent with prior literature (where increases in REM time were also reported [32]) as well as our own prior data suggesting that PPN stimulation during sleep can result in acute sleep fragmentation [19].

Upon finalization of the night-time algorithm according to the principles described in the prior session, these were tested against a night of continuous ‘daytime’ settings (24 hrs invariable gamma frequency and amplitudes, as would have been the clinical norm) (Fig. 6). We noted that having a tailored algorithm in this case of RBD resulted in a decrease in sleep fragmentation (49.4% reduction in wake epochs after sleep onset) as well as an increase in slow-wave sleep (53.1%) overall, consistent with subjective reports (blinded to settings) of the patient experiencing deeper sleep on a scheduled night-time algorithm and feeling well-rested in the morning. Therefore upon trial completion, these tailored settings were selected as ongoing therapy to ensure optimal treatment effects on daily functioning and overall quality of life.

**Fig. 6.**
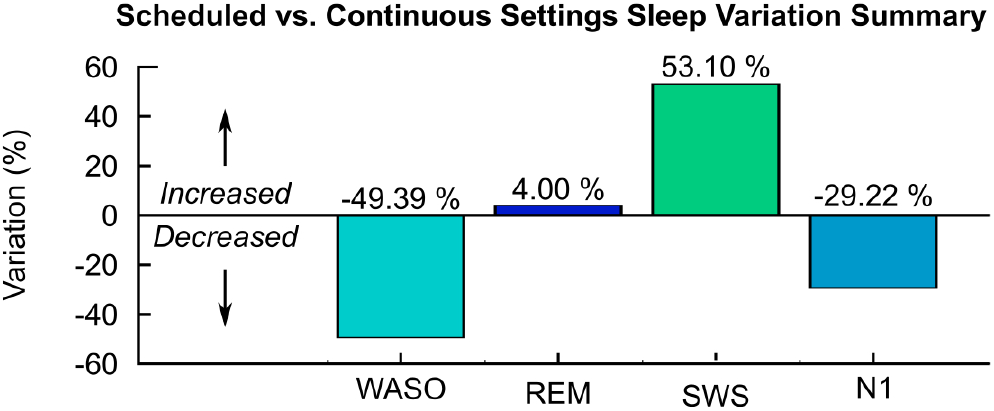
Comparison of overnight scheduled settings with continuous gamma stimulation (PSG electrophysiological summary data). Scheduled settings are associated with an increase in slow-wave sleep (SWS) and reduction in sleep fragmentation (wake after sleep onset, WASO) in the RBD phenotype. Data was recorded during the same hospital visit, with lights-off at the same time in the night and identical medication schedules. Polysomnography scored and analysed as per AASM criteria.

## IV. Discussion

We describe a novel concept of targeted neurostimulation of subcortical networks crucial for the regulation of sleep and wakefulness, aimed at normalizing diurnal and circadian rhythms in the context of neurodegenerative disorders. Representative clinical data from our ongoing trial further illustrate the applicability of these concepts. This is preliminarily evidenced by the distributed fronto-cortical network activity modulation observed longitudinally in our patients, that is confirming engagement of ascending arousal networks. Such changes and avoidance of cortical oscillatory slowing could underlie improvement in daytime alertness and functioning. The capacity of our bioelectronic platform for targeted patient-oriented treatment in the context of disorders of sleep and wakefulness is reflected by encouraging results in the case of sleep pathology, in addition to built-in device specifications.

## V. Limitations

Currently, our evidence of bioelectronic Zeitgeber efficacy is limited due to the preliminary stages of the trial where this concept is being deployed, as well as the lack of continuous home-based electrophysiological monitoring. Such a capacity made possible by wearables as well as home-based data gathering – will also alleviate one of this study’s current constraints, which is the need for multiple in-patient visits in the post-COVID climate of severe healthcare provision pressures in the United Kingdom. Longitudinal tracking of sleep diaries and repeat polysomnography in more patients with concomitant sleep and/or wake pathology, recruited in the trial, will hopefully reveal whether there are persistent beneficial effects of this new, diurnally weighted stimulation strategy. We further hope that further longitudinal data will also provide additional support that this system can successfully entrain patients’ rhythms towards a desired objective and perhaps also beneficially impact disorder symptomatology by long-lasting modifications at a circuit and perhaps even cellular level.

## VI. Conclusions and Future Steps

There is evidence that sleep disturbance exacerbates neuroinflammation in the context of neurodegeneration [33], therefore reduction of sleep fragmentation may modify a predisposing factor to worse disease progression. Encouraging effects on cognitive function after the restoration of more physiological patterns of sleep architecture have been demonstrated in in vivo animal models [34]. Addressing neuronal circuit degradation in the human, though restoration of circadian processes specifically delivered by brain-machine interfaces restoration of circadian processes is a very attractive concept. However such evidence of a bioelectronic Zeitgeber model impact in the human is (to our knowledge) yet to be discovered. An interesting follow-up point in this vein, requiring a large longitudinal cohort, would also be the comparison of both motor and autonomic outcomes between those with optimal neurophysiological and behavioural circadian responses to this experimental stimulation treatment and non-responders to circadian and diurnal re-alignment.

As a further advancement of the current stimulation algorithm, in order to increase efficacy in reducing daytime sleepiness symptoms and preserve sleep integrity, would be the development of closed-loop patterns based on local subcortical network activity or state-specific cortical patterns. For instance, the capacity to trigger stimulation while sensing oscillatory activity that corresponds drowsiness or in response to the onset of EEG slowing could facilitate alertness optimization on a fine-grained temporal scale. Such finer-tuned interventions would be valuable in patients with neurodegerenative disorders but perhaps of even higher significance in the case of those suffering from disorders of consciousness, where quality of life is severely impaired and rhythm disruptions follow subtler patterns, while also not being as easily linked to diurnal changes in behaviour. The latter cluster of conditions is therefore an active area of interest of our broader collaborative efforts.

## Acknowledgments

We would like to thank our patients and their carers for their trust, time and patience. We thank Mr Sean Martin, Dr Nagaraja Sarangmat and the Neurosciences Inpatient Service team for patient care and recruitment in the MINDS trial. Last but not least, we are grateful towards Bioinduction Ltd (particularly Hannah Campbell, Guy Lamb, and Tara Noone) for technical support during this project.

## Data Availability

The authors will consider requests to access the data in a trusted research environment. Contact.

